# Mapping the Central and Peripheral Projections of Lung Innervating Sensory Neurons

**DOI:** 10.1101/2021.05.05.442702

**Authors:** Yujuan Su, Justinn Barr, Abigail Jaquish, Jinhao Xu, Jamie M Verheyden, Xin Sun

## Abstract

While best known as the gas exchange organ, the lung is also critical for sensing and responding to the aerosol environment in part through interaction with the nervous system. The rich diversity of lung innervating neurons remains poorly understood. Here, we interrogated the cell body location, projection pattern and targets of lung-innervating sensory neurons. Retrograde tracing from the lung labeled neurons primarily in the vagal ganglia, in a spatially distributed population expressing markers including *Vglut2, Trpv1, Tac1, Calb1* or *Piezo2*. Centrally, they project to the nucleus of the solitary tract in the brainstem. Peripherally, they project along the branching airways and terminate on airway smooth muscles, vasculature including lymphatics, and selected alveoli. Notably, a discrete population of Calb1+ neurons preferentially innervate pulmonary neuroendocrine cells, a demonstrated airway sensor population. This comprehensive illustration of the properties of lung innervating sensory neurons serves as a foundation for understanding their function in lung.

## Introduction

The lung is richly innervated with nerves^1–5^. Growing evidence demonstrates that these nerves serve critical functions. Vagus nerves that innervate the airways are essential for airway protective responses, including bronchospasm and cough reflex^6–9^. Recent studies have shown that a subset of vagal afferent neurons control breathing at birth and during homeostasis^10, 11^. Sensory nerves also regulate airway hyperreactivity and immune responses^12–14^. Despite demonstrated importance, there are many open questions including the projection pattern and target specificity of lung innervating neurons, and how sensory signals from the lung are integrated by the central nervous system and translated into efferent outputs either back to lung or to other tissues.

Like the lung, many if not all internal organs are innervated. Increasing evidence demonstrates that tissue-nervous system interaction plays critical roles in physiology. A recent swell of findings led to resurgence of the term “interoception” to define the process by which the nervous system senses and integrates information from tissues^15^. For the lung, there is increasing evidence that interoception is a critical aspect of lung biology.

Nerves in the lung can be separated into sensory afferent nerves versus parasympathetic or sympathetic efferent nerves^5, 16–18^. Both afferent and sympathetic efferent nerves originate from neurons extrinsic to the lung, while parasympathetic efferent nerves can originate from neurons either intrinsic or extrinsic to the lung. Among these lung innervating neurons, afferent neurons have arguably the most documented diversity. They reside in either the vagal ganglia (VG) or the dorsal root ganglia (DRG), each composed of genetically diverse groups of sensory neurons^19–24^. Some of the diversity may stem from different developmental origin. For example, in mouse, the VG is composed of a fusion of the nodose ganglia which arise from epibranchial placodes, and the jugular ganglia which arise from neural crest^25^.

The lung is mainly innervated by nodose neurons, while the trachea and larynx are mainly innervated by jugular neurons^26–29^. Recently, manually picked vagal neurons that were retrogradely labeled from the trachea/bronchi were individually sequenced to yield the first expression profile of vagal neurons that innervate extrapulmonary airway^30^.

Vagal sensory neurons are pseudounipolar, with each cell body sending out a single axon that divides into one central- and one peripheral-projecting process^8^. Collectively, VG neurons centrally project to multiple regions of the medulla portion of the brainstem, including the nucleus of the solitary tract (nTS), the paratrigeminal nucleus (Pa5), the spinal trigeminal nucleus (Sp5), the area postrema (AP), passing through the solitary tract (TS) region^31–33^. Detailed mapping has shown that nodose neurons are part of the visceral ascending circuit that projects to nTS, while jugular neurons are part of the somatic ascending circuit that projects to Pa5^31, 32^. Vagal neurons with different molecular signature such as Trpv1 and Tac1 show overlapping but also distinct brainstem targets^33^.

In this study, we focus on mapping the central and peripheral projections of lung innervating sensory neurons. Consistent with previous reports, we found that lung innervating sensory neurons reside primarily in the VG, with a minor contribution from the DRG^24, 27, 33^. Using *in situ* hybridization combined with retrograde tracing from the lung, we found that lung innervating neurons express signature genes in a subset of vagal neuron clusters. Centrally, lung innervating sensory neurons project to the nTS region of the brainstem. Peripherally, lung innervating sensory neurons project along the airways to terminate in bronchi/bronchioles, vasculature/lymphatics, alveoli and pulmonary neuroendocrine cells (PNECs). Using several vagal subtype signature cre lines, we mapped their shared and distinct projection patterns and targets in the lung. The distinct projection patterns of these subtypes both in the brainstem and in the lung suggest possible diverse roles of these neurons in lung interoception.

### Results

#### Lung innervating sensory neurons reside primarily in vagal ganglia and project to the nTS region of the brainstem

To determine the location of lung innervating neurons and their projection paths, we genetically labeled nerves by intratracheal instillation (i.t.) of retrograde rAAV2-retro-cre virus into *Rosa-lxl-tdTomato* (*Ai14*) mice (Fig.1A). To ensure labeling specificity, all tissues were examined for fluorescence and only samples with lung-specific labeling without spillage into other organs were used for further analysis (Supplementary Fig.1A). For lung innervating sensory neurons, we examined both VG and DRG and found that compared to maximally ~3-5 labeled cells in the DRG, ~30-100 cells (a number proportional to the extent of lung infection) were labeled in the VG (Fig.1B). CUBIC (Clear, Unobstructed Brain/Body Imaging Cocktails and Computational analysis protocol) tissue clearing^34^ followed by light sheet imaging showed that vagal neuron cell bodies as well as both central- and peripheral-projecting processes were labelled (Fig.1B-D, Supplementary Movie 1). Within the VG, both large and small cell bodies were labeled, indicating that lung innervating vagal neurons are heterogeneous. We confirmed these results from rAAV2-retro-cre by using i.t. administration of several other reagents, including rAAV2-retro-GFP, fast blue and wheat germ agglutinin 594 (WGA594) in wild-type mice (Supplementary Fig.1B-D). These findings together indicate that lung innervating sensory neurons reside primarily in the VG, are of varying sizes and are scattered across VG rather than localized in a specific domain.

**Figure 1.**
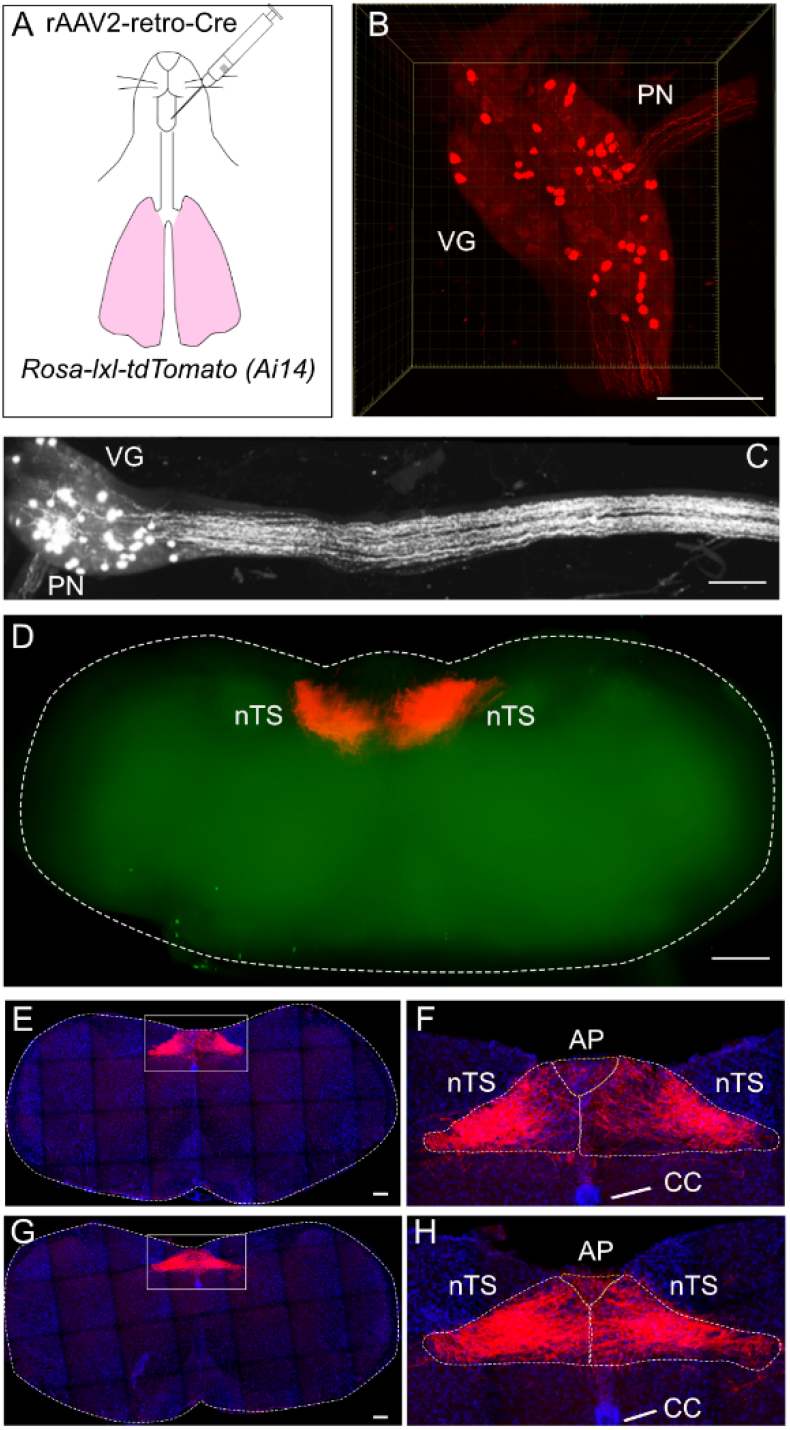
Lung innervating sensory cell bodies reside in the vagal ganglia and project to the nTS. **(A)** Intratracheal instillation scheme. **(B, C)** CUBIC cleared vagal ganglia (VG) and nerve tracts from *Rosa-lxl-tdTom* (*Ai14*) mice with intratracheal instillation of rAAV-retro-cre. PN, pharyngeal nerve. Thick nerve tract in C projects to lung. **(D)** CUBIC cleared whole brainstem of *Ai14* mice with intratracheal instillation of rAAV2-retro-cre, showing nerve terminal signals in bilateral nTS. **(E-H)** Sections (40um) of brainstem medulla at either rostral (E, F) or caudal (G, H) levels showing nerve signals. Dashed circles outline the brainstem. Boxed regions in E and G are magnified in F and H, respectively. nTS, nucleus of the solitary tract; AP, area postrema; CC, central canal. Scale bars 250um in B, 100um in C, 500um in D, 50um in E and G.

To map central projections of lung innervating sensory neurons, we cleared the whole brainstem following i.t. of rAAV2-retro-cre virus into *Ai14* mice. As shown in Fig.1D and Supplementary Fig.2, fibers projecting from lung innervating sensory neurons densely arborized in the nTS bilaterally, with a few fibers terminating in the AP. To confirm this pattern, we serially sectioned the whole brainstem. According to the mouse stereotaxic map^35^ and consistent with our cleared, light sheet image of the whole brainstem, the lung innervating vagal sensory neurons mainly project to the ventromedial nTS at both rostral (Fig.1E, F) and caudal (Fig.1G, H) levels, with only sparse projections to the AP region.

#### Vagal sensory nerves split into multiple paths to innervate the trachea or the lung

*Vglut2*, encoding a vesicular glutamate transporter, is reported to be expressed by a majority of vagal sensory neurons^10^. This is demonstrated in the confocal z-stack projection of VG from *Vglut2-ires-cre; Ai14* mice, where abundant expression of tdTomato labeled vagal neurons of different sizes (Fig.2A, B). We carefully dissected the tissue to preserve the VG, the vagal nerve trunk and all nerve branches connected to the trachea and lung. While there is sample-to-sample variation, in general the main tdTomato+ vagal sensory nerve trunk splits into multiple branches, including the glossopharyngeal nerve (GN), the pharyngeal nerve (PN), the superior laryngeal nerve (SLN) and the recurrent laryngeal nerve (RLN) branches (Fig. 2C-G, Supplementary Fig.3). These branches innervate the pharynx, the larynx and the proximal versus distal trachea, while the main vagal nerve trunk follows along the dorsal-medial side of the bronchi to project deep into the lung (Fig.2H). We repeatedly found that the PN branch is observed only on the left, but not the right side of the trachea. While it remains possible that this is a dissection artifact, it may reflect an anatomy difference between the left and right sides.

**Figure 2.**
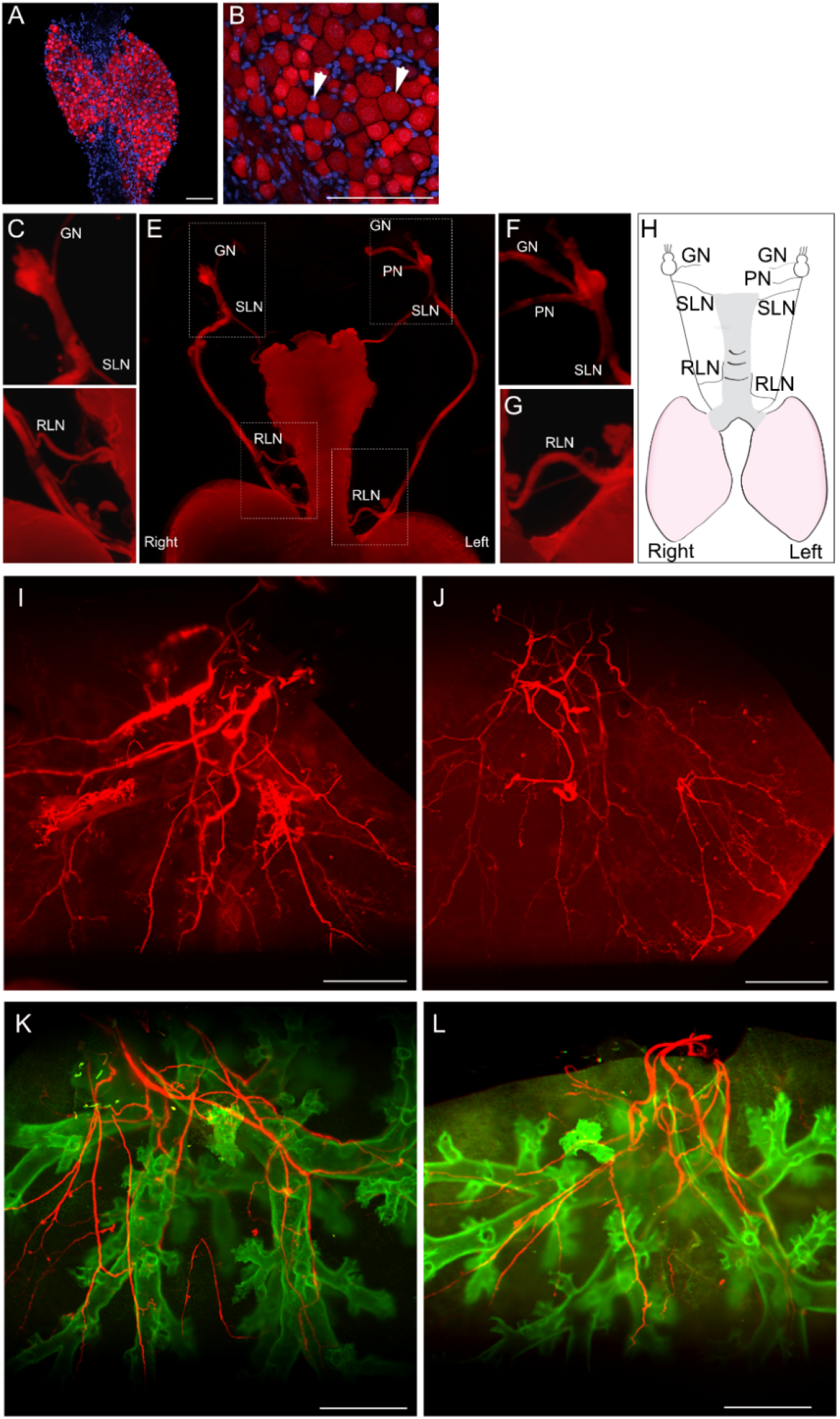
Sensory innervation pattern of the respiratory tract and lung. **(A, B)** Confocal max projection of *Vglut2-cre; Ai14* vagal ganglia. tdTomato labels both large and small neurons (arrowheads in B). **(C-G)** Representative *Vglut2-cre; Ai14* respiratory tract with vagal ganglia and nerve bundles. **(H)** Diagram depicting the consensus pattern of respiratory tract innervation by vagal sensory nerves. **(I, J)** Light sheet images of *Vglut2-cre; Ai14* whole cranial lung lobes from two mice at similar age. GN, glossopharyngeal nerve; PN, pharyngeal nerve; SLN, superior laryngeal nerve; RLN, recurrent laryngeal nerve. **(K, L)** Light sheet images of *Nkx2-1^GFP^; Vglut2-cre; Ai14* whole cranial lung lobes from two mice at similar age. Left and right side of the mouse is labeled in E and H. Scale bars 100um in A, B, 1mm in E, I-L.

To comprehensively depict sensory nerves in lung, we cleared whole cranial lobes of *Vglut2-ires-cre; Ai14* lungs using CUBIC and documented tdTomato+ nerve patterns using light sheet imaging (Fig.2I, J, Supplementary Movie 2). In all samples, multiple thick nerve bundles enter the lung at the proximal bronchus. As they project distally, they split into thinner fibers. To better track the nerve path, we introduced *Nkx2-1^GFP^* to outline the airway epithelium^36^ in the *Vglut2-ires-cre; Ai14* background. Whole lung lobe imaging showed that Vglut2+ sensory nerves project and branch along with the airways (Fig.2K, L, Supplementary Movie 3). In contrast to the stereotyped branching pattern of airways^37^, the innervation pattern of nerve fibers within the lung is more variable even among equivalent lobes of adult mice of similar age (Fig.2I-N). A few fibers project all the way to the periphery of the lung while most terminate as thin threads at the end of distal airways. Knob-like sensory nerve endings can be detected on occasion, but the position of these elaborate endings is not stereotyped (Fig.2I and Supplemental Movie 2).

#### Molecular signature of lung innervating vagal sensory neurons

Vagal sensory neurons are known to be heterogeneous by multiple measures, including developmental origin, cell size, conduction velocity and molecular characteristics^19, 22^. To define the molecular signature of lung innervating neurons, we used RNAscope *in situ* hybridization assay to probe expression of tdTomato+ neurons in the VG after i.t. of rAAV2-retro-tdTomato virus into the lungs of wild-type mice. Based on recently published VG single-cell RNA sequencing (scRNA-seq) data^19, 21, 30^, we identified 12 genes that either individually or in combination define all vagal neuron types. Two-color RNAscope *in situ* hybridization analysis revealed that as expected, all *tdTomato+* neurons express *Vglut2*, a marker expressed in a majority of neurons. Similarly, a high proportion of *tdTomato+* neurons express nociceptor *Trpv1*. In comparison, a lower proportion of *tdTomato+* neurons express *Tac1, Calb1, Piezo1, Piezo2, Trpa1* and *Runx3*, while only a few lung innervating neurons express *Vip* and *Tmc3*, and almost none expressed *Glp1r* or *Gabra1* (Fig.3A-L).

**Figure 3.**
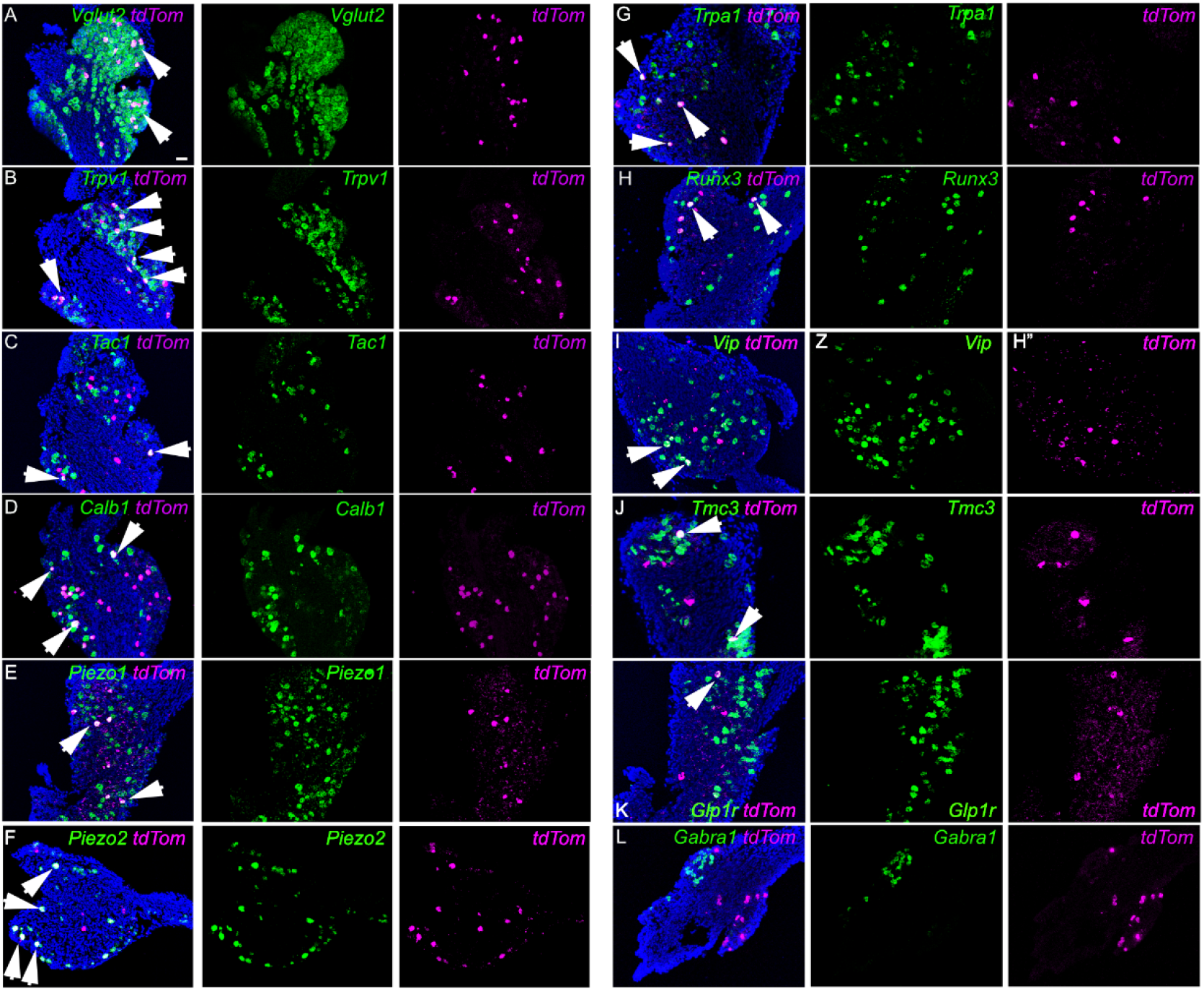
Transcriptional signature of lung innervating vagal neurons. **(A-L)** Lung innervating vagal neurons are retrogradely labeled with rAAV-retro-cre into *Ai14* lungs (*tdTom*). RNAscope was performed on vagal ganglia sections for overlap between *tdTom* (magenta) and *Vglut2, Trpv1, Tac1, Calb1, Piezo1, Piezo2, Trpa1, Runx3, Vip, Tmc3, Glp1r* and *Gabra1* (green), individually as labeled. Arrowheads point to neurons with double labeling. Scale bar 50um.

#### Different vagal sensory neuron subgroups show distinct central projection targets in the medulla of brainstem

To determine if vagal neurons with distinct molecular signatures may exhibit distinct central and peripheral projection patterns, we selected several representative markers to investigate further. We utilized *Vglut2* as a global marker of sensory neurons, *Trpv1* as a nociceptor receptor marker, *Tac1* as a neuropeptidergic neuron marker and *Calb1* as a large-diameter neuron marker. Double RNAscope staining showed that *Calb1* expression exhibited only minor or no overlap with either *Trpv1* or *Tac1* expression (Supplementary Fig.4A-F). Between *Trpv1* and *Tac1* patterns, based on published immunostaining^33^, scRNA-seq data^19^, and confirmed by our staining, *Trpv1* and *Tac1* are co-expressed in the jugular neurons, but not nodose neurons (Supplementary Fig.3G-I, Supplementary Fig.4J-L). Jugular neurons are known to innervate the trachea^26–29^ while our lung-originated retrograde tracing labeled primarily nodose neurons (Fig.1B and C, Supplementary Fig.1B-D). And in nodose neurons, *Trpv1* and *Tac1* represent two largely distinct populations.

To study innervation patterns of these selected vagal neuron subtypes, we injected cre-dependent AAV-flex-tdTomato directly to unilateral VG of either *Vglut2-ires-cre, Trpv1-ires-cre, Tac1-ires-cre or Calb1-ires-cre* mice (hereafter referred to as marker-cre lines) (Fig.4A). Three weeks after injection, tdTomato fluorescence is observed in vagal neuron cell bodies (Fig.4B-E). As expected, among the four lines, most labeling was observed in *Vglut2-cre* and least labeling was observed in *Calb1-cre*.

**Fig. 4.**
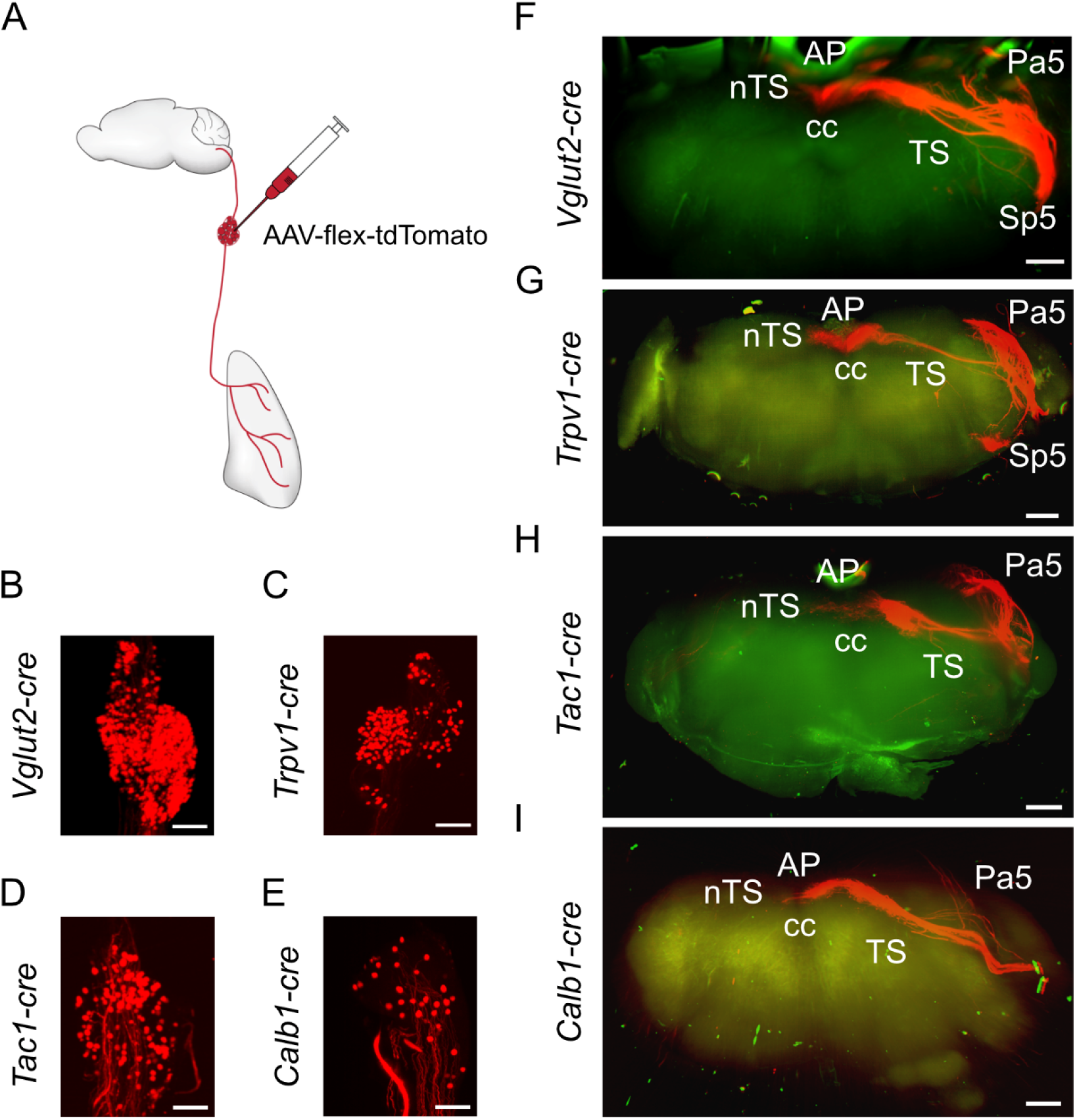
Vagal neuron labeling and brainstem projection of *Vglut2-cre, Trpv1-cre, Tac1-cre and Calb1-cre* mice after unilateral vagal ganglia injection of AAV-flex-tdTomato. **(A)** Schematic for AAV-flex-tdTomato vagal microinjection. **(B-I)** Representative CUBIC cleared vagal ganglia (B-E) and whole brainstem (F-I) after AAV-flex-tdTomato vagal microinjection to the respective cre lines as labeled. Autofluorescence green channel was used as background to outline the brainstem (F-I). nTS, the nucleus of solitary tract; TS, the solitary tract; AP, the area postrema; Pa5, the paratrigeminal nucleus; Sp5, the spinal trigeminal nucleus, CC, central canal. Scale bars 200um in (B-E), 500um in (F-I).

To clearly visualize the central projection pattern of these vagal sensory neuron types, we cleared the whole brainstem using CUBIC and performed light sheet imaging. Collectively, the axons of vagal sensory neurons are known to be present in nTS, AP, TS, Pa5 and SP5 regions in the medulla^31–33^. Consistent with this, our mapping using pan-VG line *Vglut2-cre* labeled vagal fibers are found in nTS, TS, Pa5, SP5, and to a limited extent in AP (Fig.4F, n=5 animals). While there is an enrichment of signals ipsilateral to the infected ganglia, arborizations were also observed in the contralateral nTS. Consistent with a recent report^33^, *Trpv1-cre* labeled fibers were found primarily in bilateral nTS and ipsilateral TS, Pa5, with minor arborizations in AP or SP5 (Fig.4G, n=6 animals). Most *Tac1-cre* labeled afferents were found in the ipsilateral nTS, TS and Pa5 regions, with a few contralateral nTS terminating fibers (Fig.4H, n=5 animals). For *Calb1-cre* labeled vagal neurons, fibers were mainly concentrated in the ipsilateral nTS and TS, with little innervation of other medulla regions (Fig.4I, n=5 animals).

Aside from these four lines, we also investigated projection pattern of Piezo2+ neurons, a key mechanosensory population in the VG^19, 21, 30^. We injected cre-dependent AAV-flex-tdTomato directly to unilateral VG of *Piezo2-ires-cre* mice. This led to labeling of neurons in the VG at variable but in general low frequencies. Interestingly, even in animals with the more abundant labeling in the VG (Supplementary Fig.5A, n=4 animals), there is only sparse central projections to Pa5 and Sp5, and minimal projections to nTS (Supplementary Fig.5B). Equally sparse peripheral projections were found in the lung (Supplementary Fig.5C, D). Due to this low frequency of labeling, we focused on the remaining four lines for further investigation in lung.

#### Vagal sensory neurons project to both airway and alveolar regions

As sensory innervation of the lung originates primarily from the VG (Fig.1B and C, Supplementary Fig.1), we expected a similar pattern of lung innervation from vagal injection of AAV-flex-tdTomato into *Vglut2-cre* animals as we did from *Vglut2-cre; Ai14* animals from a genetic cross. We found this to be the case, and each provide a comprehensive illustration of lung sensory innervation pattern (Fig.5, Supplementary Fig.6). In both cases, vagal sensory neurons project thick bundles of nerve fibers that entered the lung following along the major bronchi, and projected to both proximal, distal airways as well as the alveolar region.

To study the pattern of vagal subtypes, we performed unilateral vagal injection of cre-dependent AAV-flex-tdTomato into *Trpv1-cre, Tac1-cre* and *Calb1-cre* mice, and assessed the pattern of nerve innervation within the lung. Similar to vagal injection of *Vglut2-cre*, thick nerve bundles were observed parallel to proximal airways for *Trpv1-cre, Tac1-cre* and *Calb1-cre* labeled vagal neurons (Fig.5A-H). These bundles run in the proximal-to-distal direction and split into thin fibers in terminal bronchioles that open into the alveolar region (visualized using anti-Podoplanin, PDPN) (Fig.5I-P). Among the four lines, *Vglut2-cre* and *Trpv1-cre* labeled nerves frequently project beyond bronchioles into the alveolar region, while only in a small subset of tissues, *Tac1-cre* labeled nerves were found in the alveolar region (rare example shown in Fig.5K, L). *Calb1-cre* labeled nerves target the airway, but not the alveolar region (Fig.5O, P).

**Figure 5.**
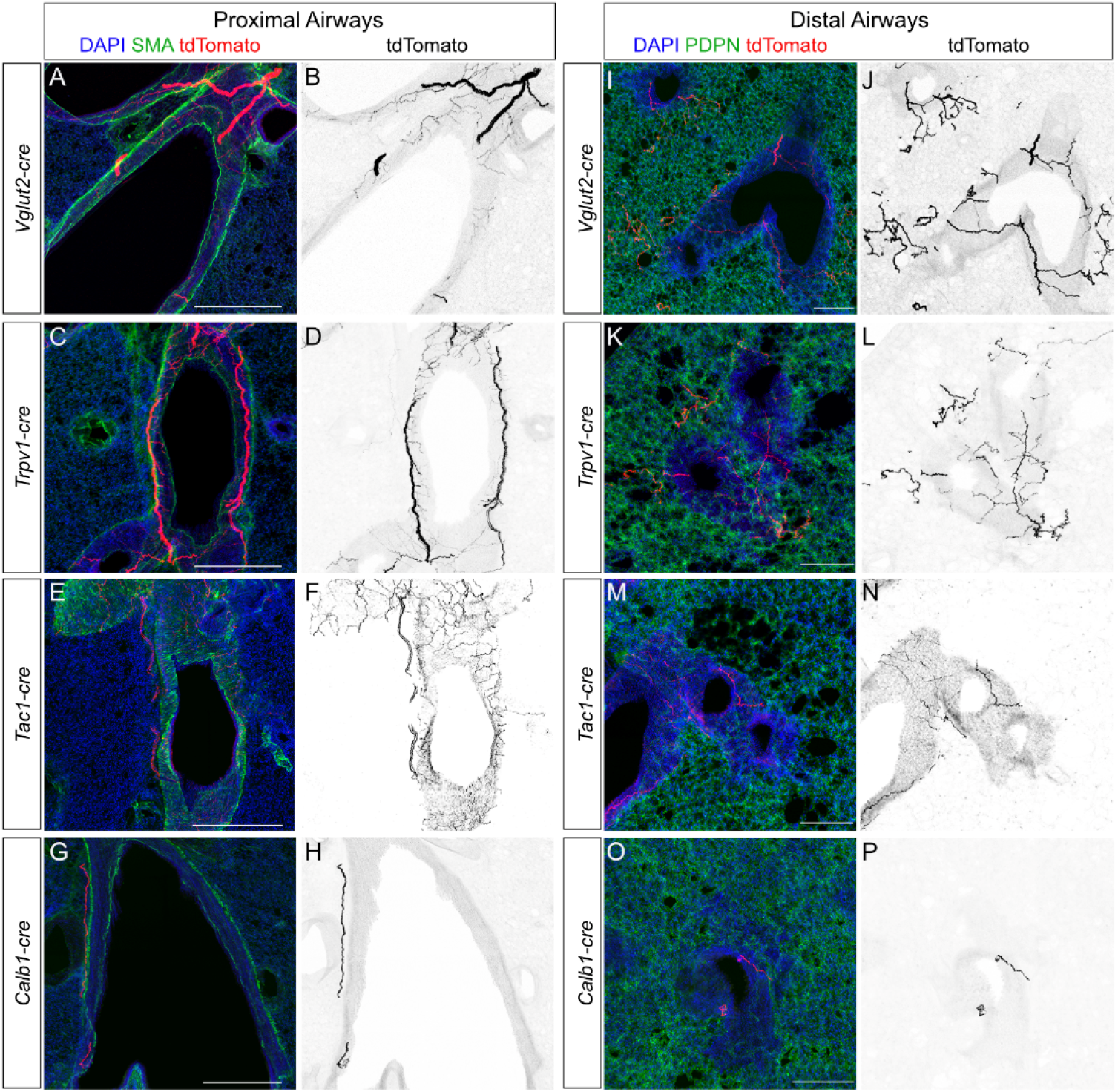
Vagal sensory neurons project to proximal and distal airways within the lung. **(A-P)** Sections (99um) of the lung showing vagal nerve projections near proximal airways (A-H) or distal airways (bronchioles, I-P) within the lung following vagal injection of AAV-flex-tdTomato into the respective cre lines as labeled. Airway smooth muscles were marked by SMA staining. The alveolar region was marked by staining of PDPN expressed by alveolar type I cells. Scale bars 500um in (A, C, E, G), 200um in (I, K, M, O).

#### Different vagal sensory neuron subtypes show distinct projection patterns on airway smooth muscle

In the proximal airway, aside from thick bundles, vagal sensory nerves in *Vglut2, Trpv1, Tac1*, but not *Calb1* lines also extend thin fibers around the airway circumference in the same orientation as airway smooth muscle cells, visualized using anti-transgelin (TAGLN/SM22) antibody staining (Fig.6A-F). Near proximal airway smooth muscle, we observed elaborately ramified nerves with knob-like terminal endings from both *Vglut2-cre; Ai14* mice from genetic cross, AAV-flex-tdTomato vagal injection into *Vglut2-cre* mice, as well as *Adcyap1-cre; Ai14* mice from genetic cross (Fig.6G-L). scRNA-seq data show that *Adcyap1* is expressed in many vagal sensory subtypes^38^. Endings with this morphology were not observed from Trpv1+, Tac1+ or Calb1+ vagal neurons.

**Figure 6.**
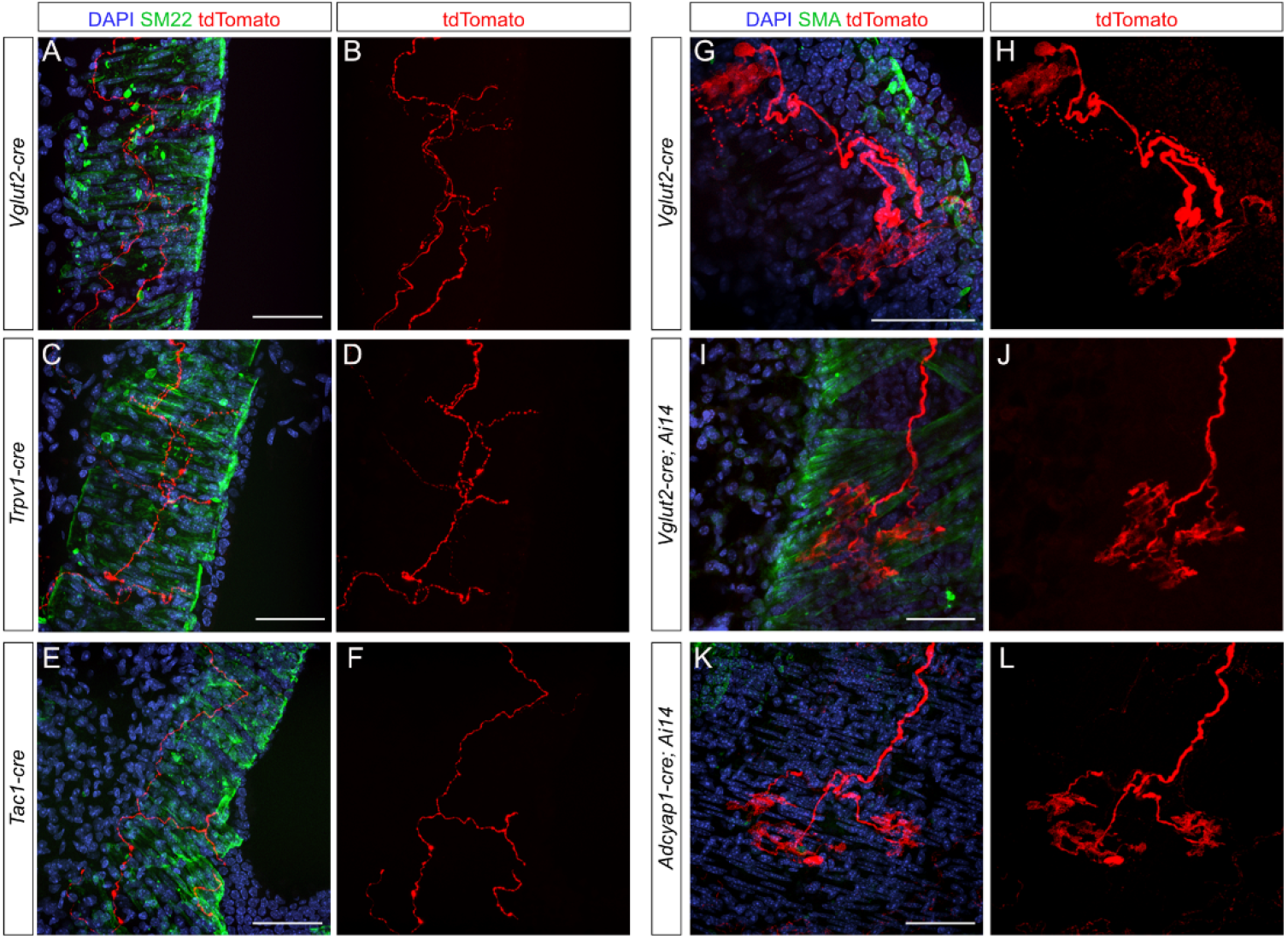
Vagal sensory neurons project to airway smooth muscles. **(A-F)** Sections (99um) of the lung showing thin nerve fibers adjacent to airway smooth muscles marked by SM22 following vagal injection of AAV-flex-tdTomato into respective cre lines as labeled. **(G-L)** Thick, branched nerve terminals near airway smooth muscle marked by SMA. This subepithelial nerve terminal morphology was observed following injection of AAV-flex-tdTomato into *Vglut2-cre* mice (G, H), in *Vglut2-cre; Ai14* mice from direct genetic corss (I, J) and in *Adcyap1-cre; Ai14* mice from direct genetic cross (K, L). All scale bars 50um.

#### Vagal sensory neurons project to selected alveolar epithelial cells

Sparse nerve terminals were found in the alveolar region, mostly near terminal airways. We investigated their targets, including alveolar type 1 (AT1, labeled with anti-receptor for advanced glycation end products, RAGE) and alveolar type 2 (AT2, labeled with anti-pro-surfactant protein C, SPC) cells. In both the *Vglut2-cre* and *Trpv1-cre* lines, rare sensory nerve projections appear to traverse between alveolar units in tortuous routes on large and flat AT1 cells (Fig.7A, B, E, F). While they ignore most AT2 cells, occasionally, the nerves pass by selected AT2 cells with increased density, suggestive of interaction (Fig.7C, D, G, H).

**Figure 7.**
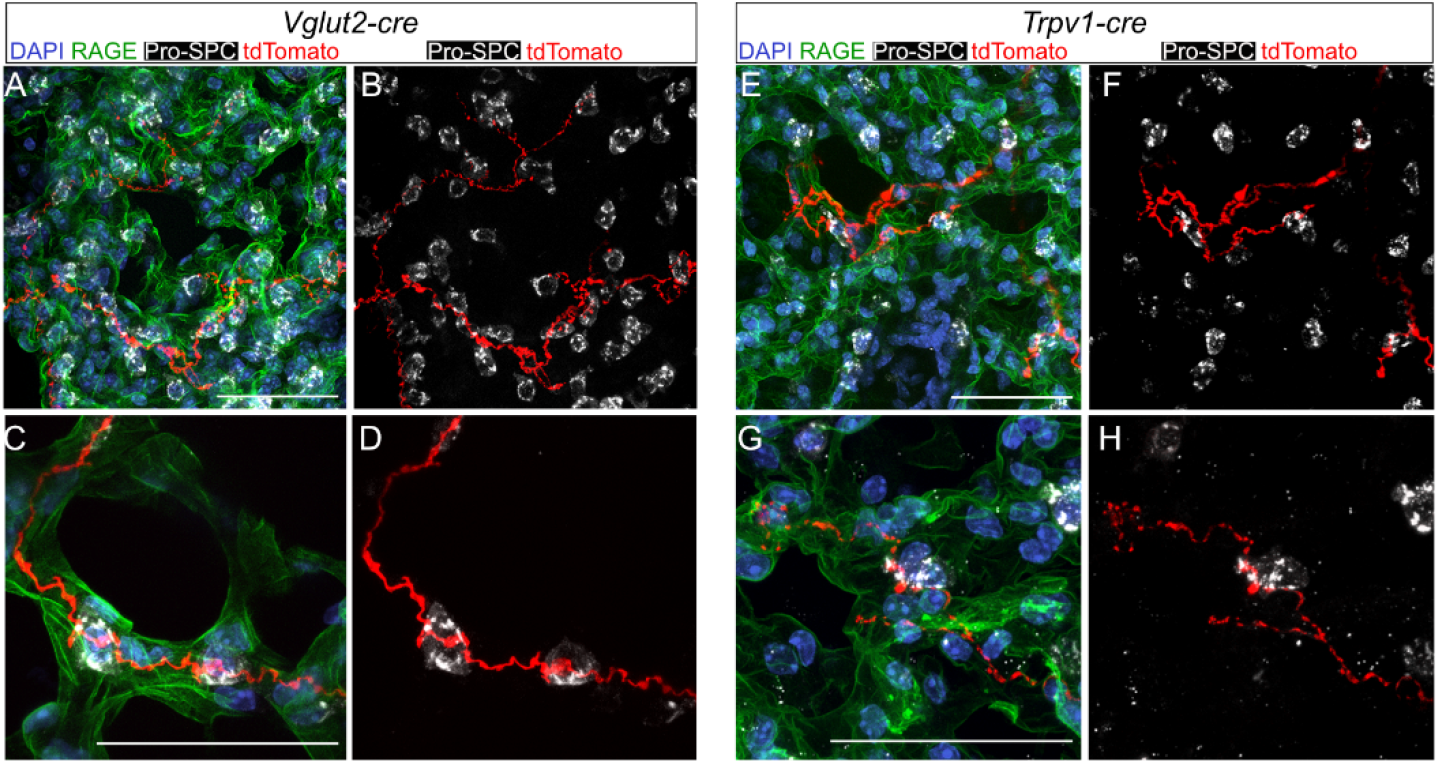
Vagal sensory neurons project to the alveolar region. **(A-H)** Sections (99um) of the alveolar region showing thin nerve fibers on selected AT1 cells, marked by RAGE, and AT2 cells, marked by Pro-SPC, following vagal injection of AAV-flex-tdTomato into *Vglut2-cre* (A-D) or *Trpv1-cre* (E-H). All scale bars 50um.

#### Vagal sensory neurons project sparsely to lung vasculature

Compared to thick vagal sensory bundles localized near airways, sparse nerve fibers were found adjacent to vasculature. Using Von Willebrand Factor (VWF) as a marker of large vessels including arteries and veins, we found that *Vglut2-cre* and *Trpv1-cre* labeled fibers project to vasculature, while only rare *Tac1-cre* and *Calb1-cre* labeled fibers were found adjacent to vasculature (Fig.8A-H). In comparison to arteries and veins, more nerve fibers were found adjacent to lymphatic vasculature marked by immunostaining for lymphatic vessel endothelial hyaluronan receptor-1 (LYVE) (Fig.8I-P). Vglut2+ and Trpv1+ sensory nerve fibers appear to interact with lymphatic vessels, while Tac1+ and Calb1+ sensory fibers appear to pass by them. A similarly sparse innervation pattern of vasculature and lymphatic vessels were observed from thick lung cryosections of *Vglut2-cre; Ai14* genetic cross (Supplementary Fig.7). In contrast, the sympathetic efferent nerves, as marked by tyrosine hydroxylase (TH) antibody, innervate the vasculature densely but the airway sparsely (Supplementary Fig.8).

**Figure 8.**
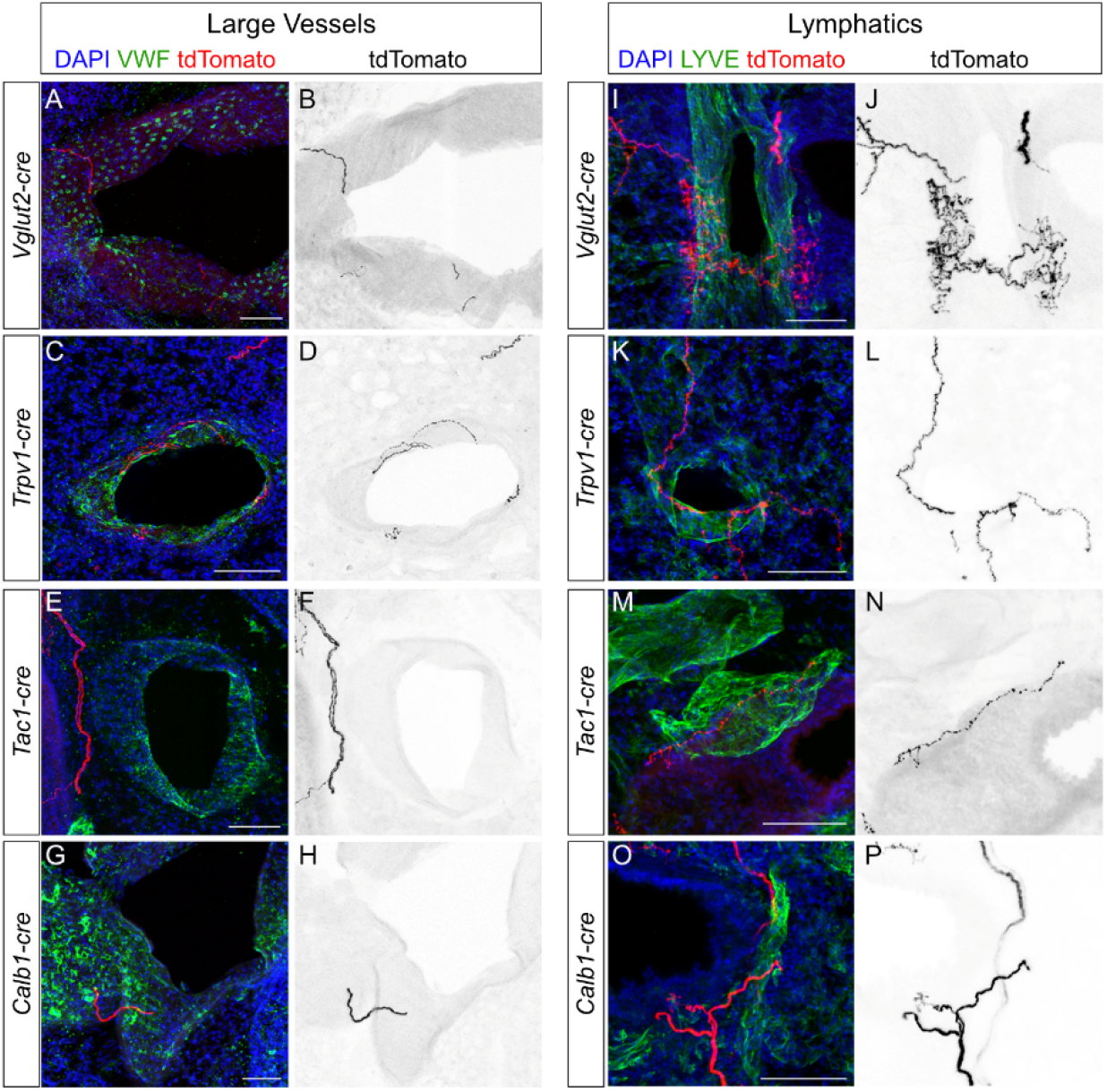
Vagal sensory neurons project to vasculature within the lung. **(A-P)** Sections (99um) of the lung showing vagal nerve projections near arteries/veins marked by VWF (A-H), or lymphatic vessels marked by LYVE (I-P) following vagal injection of AAV-flex-tdTomato into respective cre lines as labeled. All scale bars 100um.

#### *Calb1-cre* labeled nerves preferentially project to and intercalate between PNECs in NEB clusters

In mouse lung, PNECs are rare airway epithelial cells that are often present in clusters, called neuroepithelial bodies (NEBs). PNECs present dense core vesicles containing neuropeptides/neurotransmitters such as calcitonin gene-related peptide (CGRP), and are known to be innervated by both sensory and afferent nerves^23^. In *Vglut2-cre* mice injected with AAV-flex-tdTomato, labeled nerve fibers approach NEB from the basal side, and also project in-between PNECs to the apical side of the cluster (Fig.9A-C, Supplementary Fig.9A, B, Supplementary Movie 4, n=3 animals). In comparison, PNEC innervations were rarely observed from either *Trpv1-cre* or *Tac1-cre* labeled vagal nerves. But when they were found, *Trpv1-cre* labeled nerves only contact the basal side of individual PNECs, while *Tac1-cre* labeled nerves project through the NEBs (Fig.9D-I, Supplementary Fig.9C-F, Supplementary Movie 5, 6, n=4 animals each). Interestingly, we found that *Calb1-cre* labeled vagal sensory nerves, while relatively sparse in lung compared to other lines, innervate NEBs with high preference. In *Calb1-cre* animals injected with AAV-flex-tdTomato into the VG, labeled nerve fibers were frequently found to approach NEBs from the basal side, project in-between PNECs to the apical side (Fig.9J-L, Supplementary Fig.9G, H, Supplementary Movie 7, n=4 animals). As Calb1+ neurons represent a small subset of vagal neurons in published scRNA-seq data^21^ as well as our staining data (Fig.3D,4E), our findings suggest that this small subset is enriched for PNEC targeting neurons.

**Figure 9.**
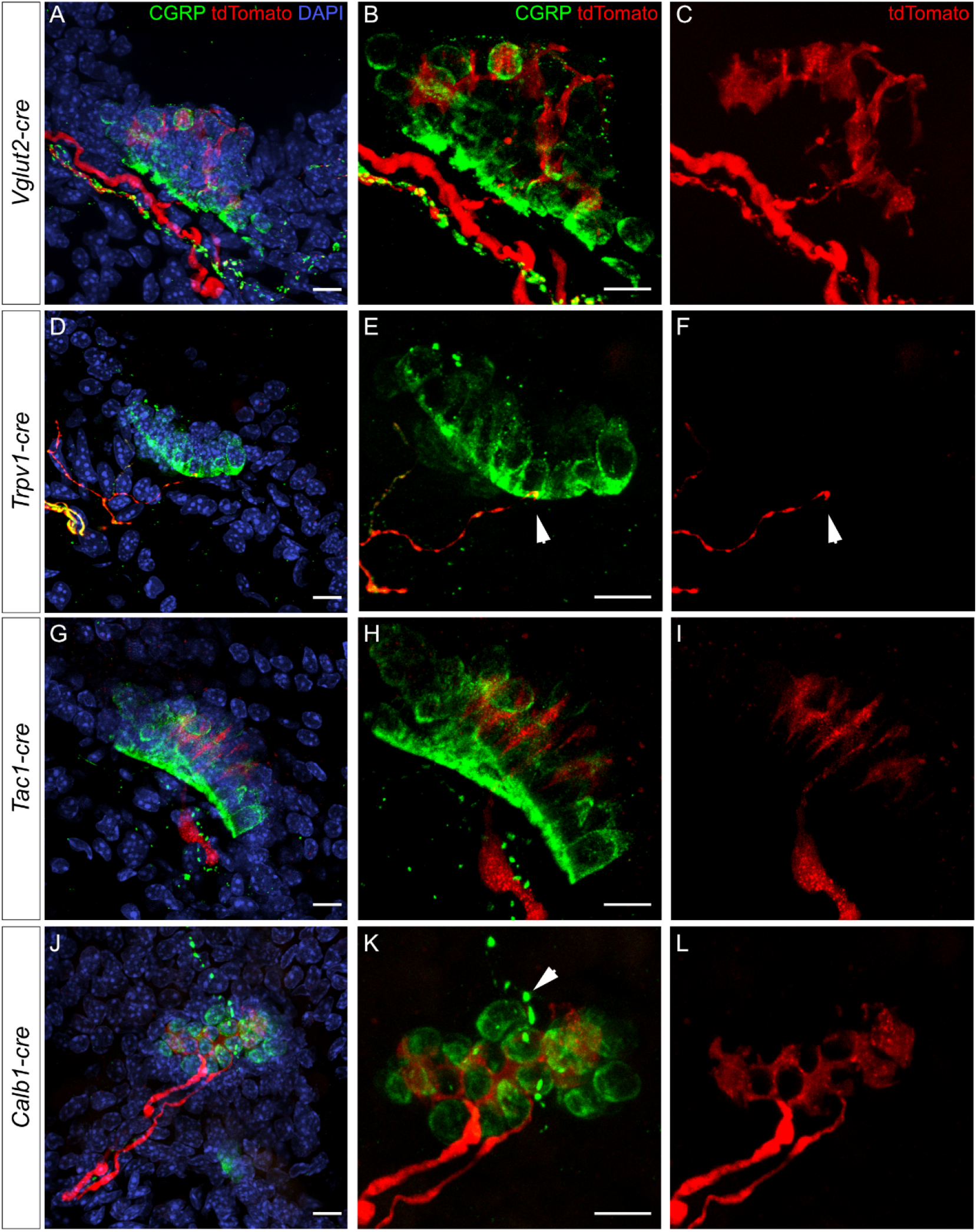
Vagal sensory neurons project to PNECs. **(A-L)** Sections (99um) of the lung showing vagal nerve projections contacting PNECs, labeled by CGRP, following vagal injection of AAV-flex-tdTomato into respective lines as labeled. *Vglut2, Tac1* and *Cabl1* projections intercalate between PNECs from the basal to apical side (B, H, K and supplemental movies 4, 6, 7). *Trpv1-cre* vagal sensory nerves only contact the basal side of PNEC clusters (arrowheads in E, F) and do not intercalate between PNECs (supplemental movie 5). Arrowhead in K indicates CGRP+ nerve fibers passing above PNEC clusters. All scale bars 10um.

### Discussion

Mapping the projection of neuronal subtypes is a critical step towards understanding neural control of organ function. Here, we found that lung innervating sensory neurons reside primarily in the VG, with a minor contribution from the DRG. Through mapping of *Trpv1-cre* labeled C-fibers, *Tac1-cre* labeled neuropeptidergic fibers and *Calb1-cre* labeled A-fibers, three representative vagal neuron types that innervate the lung, we found that they exhibit distinct projection patterns both centrally to the brainstem as well as peripherally to the lung. These results serve as the foundation for functional studies to understand their respective roles in the transmission of sensory signals from the lung to the central nervous system, in the poorly understood broader context of lung interoception^15^.

Several recent studies systematically defined transcriptomic heterogeneity of vagal sensory neurons through scRNA-seq, yielding an upward of 37 clusters^19, 21^. Retrograde tracing from the gastrointestinal tract^20^ or the trachea^30^ followed by single cell transcriptome studies started to reveal the specific signature of vagal neurons that innervate these target tissues. By RNAscope *in situ* hybridization using markers that best distinguish vagal clusters, coupled with validation using transgenic lines, our finding is consistent with prior knowledge on the types of neurons that innervate the lung^8^. For example, we observed overlapping expression between lung innervating *tdTomato+* neurons and nociceptive markers *Trpv1* and *Trpa1*. These nociceptive vagal neurons are reported to play a role in airway chemosensation and reflex control^39, 40^. We also observed some overlap between *tdTomato* and mechanoreceptors *Piezo1* and *Piezo2*. These neurons in the VG are required for baroreception, blood pressure regulation, and airway stretch-induced apnea in mice^11, 41^. It is interesting that in our still limited analysis, there is little or no overlap of lung innervating *tdTomato* signal with *Glp1r* or *Gabra1* expression. *Gabra1* is a mechanoreceptor that labels nerves with intraganglionic laminar endings that innervate the stomach^20^. *Gabra1*-expressing vagal neurons display large, isolated and branched terminals in the trachealis muscle^21^. It is possible that the *Glp1r* and *Gabra1* populations project more specific to the stomach and the trachea than to the lung. In-depth analysis of the transcriptome of the diverse lung innervating vagal neurons is a critical step towards understanding their functional heterogeneity in relaying sensory inputs from the lung.

Mapping with rAAV-retro-cre in the lungs of reporter mice showed that lung innervating neurons centrally project primarily to the nTS, with minor targeting to the AP. In comparison, gut innervating vagal neurons project more densely to the AP than lung innervating vagal neurons, aside from shared targeting to the nTS^42^. Previous retrograde tracing from the trachea or the lung separately using trans-synaptic herpes simplex virus showed that trachea innervating neurons reside in the jugular ganglia and project to Pa5, while lung innervating neurons reside in the nodose ganglia and project to the nTS^31, 32, 43^ We reasoned that no signal in Pa5 was observed in our experiment because intratracheal administration of rAAV-retro-cre delivered virus primarily to the lung with minimal infection of the trachea. It has been further suggested that vagal projection to Pa5 mediate reflex response while nodose projection to nTS mediate autonomic function^8, 31^. Among the multiple brainstem regions targeted by vagal neurons, the finding that the lung innervating neurons only target a subset suggests a mechanism that underly tissue specificity of interoception.

Our mapping delineates cellular targets and terminal morphologies of sensory nerves in lung (Fig.10). Airway smooth muscle is a major target. There, nerves display either thin free ending following *Trpv1-cre* or *Tac1-cre* labeling, or large, branched terminals following non-discriminatory *Vglut2-cre* or *Adcyap1-cre* labeling. The latter morphology is reminiscent of branched nerves with knob-like terminals that have been described in the lung and the stomach, which can be activated by stretch^44–46^. It will be interesting to determine if aside from sensing stretch during normal inhalation, the same type of nerves would respond to pathological change of mechanical cues such as in allergen-induced airway constriction in asthma.

**Figure 10.**
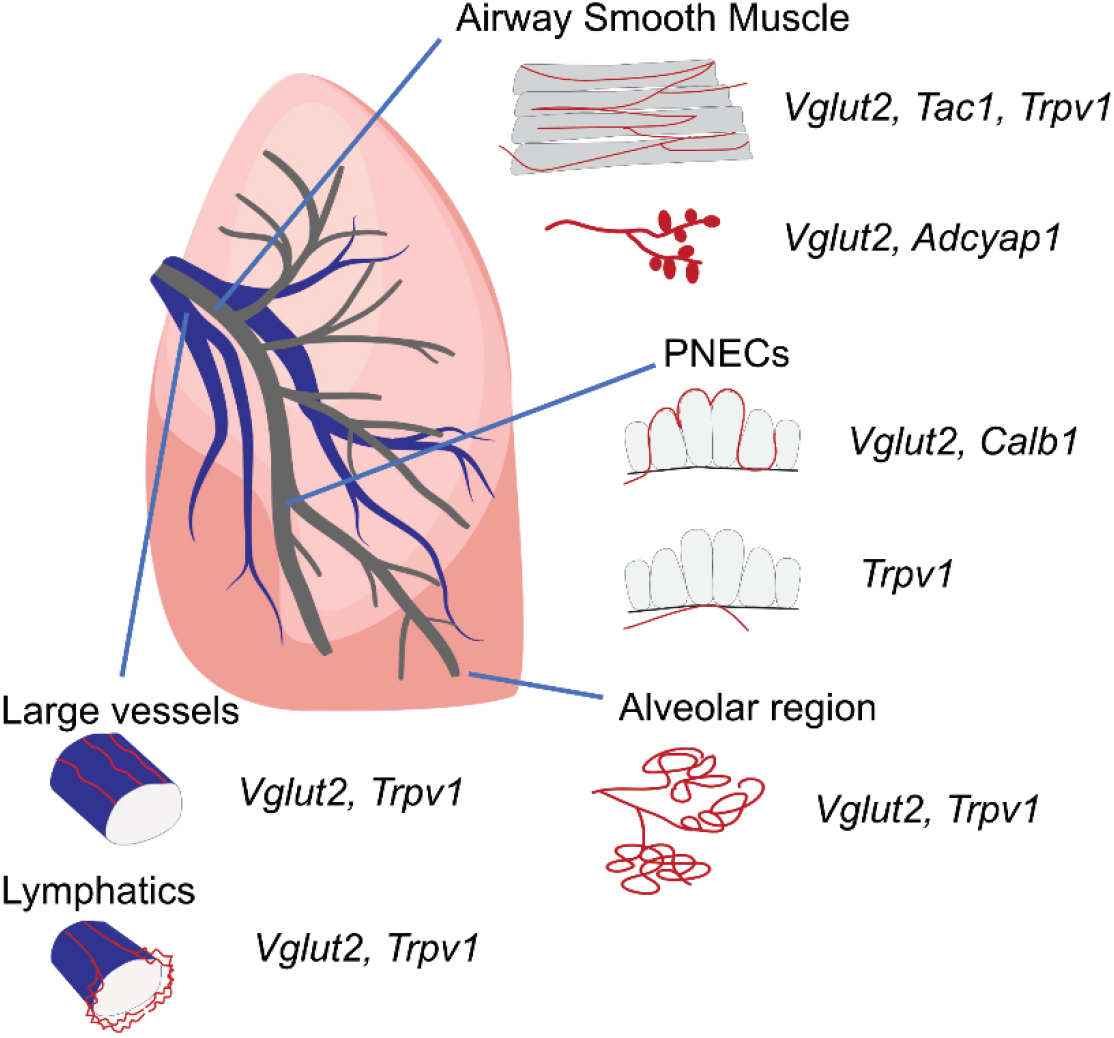
Summary of Vglut2, Trpv1, Tac1 and Calb1 vagal sensory neuron projection patterns in lung. Within the lung, *Vglut2-cre, Tac1-cre* and *Trpv1-cre* labeled thin nerve fibers on airway smooth muscles. *Vglut2-cre* and *Adcyap1-cre* labeled nerves with branched and knob-like terminals on airway smooth muscles. *Vglut2-cre* and *Calb1-cre* labeled nerve fibers that innervate the epithelium at PNEC clusters, intercalating between PNECs. *Trpv1-cre* labeled nerve fibers contact the basal side of PNEC clusters. *Vglut2-cre* and *Trpv1-cre* labeled nerves were found in the alveolar region near junction with distal airways. A sparse density of *Vglut2-cre* and *Trpv1-cre* labeled nerve fibers project to vasculature including large vessels and lymphatics.

Compared to airway innervation, we observed sparse projection of vagal afferent nerves to vasculature. Vagal sensory neurons are known to sense blood pressure from the aortic arch of the heart^41, 47^ and nutrients from the portal vein^20, 48^. Our finding is consistent with previous data in guinea pig that Trpv1+ nerves localize near vasculature^49^ and capsaicin-sensitive nerves regulate pulmonary vascular permeability^50^. Our observation that *Trpv1-cre* labeled nerves interact with lymphatics suggests that they may contribute to lymphatics function in immune cell trafficking and fluid drainage. Indeed, *Trpv1*-expressing nerves have also been shown to regulate immune responses in the lung, including to suppress the number of lung-resident gamma-delta T cells and the recruitment of neutrophils during bacterial infection^51^.

Our detection of sparse *Trpv1-cre* labeled nerve endings in the alveolar region suggest that these nociceptor-expressing nerves could sense insults such as toxins and pathogens that reach the alveoli. It is unclear if the selected AT2 cells with intense sensory nerve density are different from surrounding AT2 cells. If nerve innervation plays a role in alveolar injury repair, these AT2 cells that interact with nerves may serve as active progenitors. Alternatively, these AT2s may serve as either anchor for nerves, or scattered sensors that relay aerosol signals to nerves.

Despite their rarity, PNECs are frequent targets of vagal sensory nerves. They are airway sensors that are activated following allergen challenge^52, 53^, mechanical stretch^54, 55^ and hypoxia^56, 57^ Our labeling with *Trpv1-cre* confirmed previous finding in the rat lung that these nerves contact the basal surface of NEBs^58, 59^ In contrast, *Tac1-cre* labeled nerves intercalate between PNECs. More frequently than in either *Trpv1-cre* or *Tac1-cre*, we found that *Calb1*-expressing vagal neurons, a relatively minor subset of vagal neurons, preferentially target PNEC clusters within the lung. These neurons have large cell bodies, characteristic of myelinated A-fibers. This is consistent with previous findings from antibody staining that sensory nerve fibers that intercalate through PNECs are myelinated A-fibers^59, 60^ It awaits to be determined what signals do these *Calb1-cre* labeled neurons transmit, and how they function together with PNECs.

The lung, with its vast surface area and continuous breathing movements, serves as a prominent sensory organ for chemical and mechanical inputs. This study integrates and advances knowledge on shared and specific properties of lung innervating sensory neurons. We hope that findings here will serve as a useful reference for future studies of lung interaction with the nervous system.

### Methods

#### Animals

All experimental procedures were performed in the American Association for Accreditation of Laboratory Animal Care (AAALAC)-certified laboratory animal facility at the University of California San Diego, following protocols approved by institutional animal care and use committee (IACUC). *Vglut2-ires-cre* (028863), *Trpv1-ires-cre* (017769), *Tac1-ires-cre* (021877), *Calb1-ires-cre* (028532), *Adcyap1-2A-cre* (030155) and *Rosa-lxl-tdTomato (Ai14*, 007914) were purchased from the Jackson lab. All cre driver lines used are viable and fertile and no abnormal phenotypes were detected. *Nkx2-1^GFP^* strain was a gift from Dr. D. Kotton and Dr. L. Ikonomou and has been previously described^36^.

#### Intratracheal instillation

Mice were anesthetized with a mixture of ketamine (100mg/kg) and xylazine (10mg/kg) via intraperitoneal injection. The mouse intubation kit (Braintree Scientific) was employed. Briefly, the mouse was positioned on the surgical platform. The tongue was gently moved to the side using blunt forceps and the laryngoscope positioned for direct visualization of the glottis. Trachea was intubated with a 22-gauge catheter. For intratracheal instillation, 20ul rAAV2-retro-cre, rAAV2-retro-GFP (Boston Children’s Hospital Core, 1.5-2×10^13^ genome copies/ml, 1:10 diluted in PBS), Fast Blue (Polysciences, 2% w/v in sterile water) or WGA594 (Thermo Fisher, 5mg/mL in sterile water) were delivered. Animals recovered from the procedure were returned to housing. For analysis, mice were sacrificed 1 week after Fast Blue and WGA594 instillation, and 3 weeks after virus instillation.

#### *Vglut2-ires-cre; Ai14* mice blunt dissection

*Vglut2-ires-cre; Ai14* mice were euthanized, transcardinally perfused with PBS to remove blood. Using fine forceps, bilateral VG and the vagal nerve trunk were isolated with all branch connections preserved. All tissues were moved to a large petri dish filled with PBS and imaged under bright field or fluorescence using an Olympus MVX-10 microscope.

#### Vagal injection

Under anesthesia, unilateral vagal complex of adult mice was surgically exposed by making an incision along the ventral surface of the neck. A micropipette containing 200nl AAV2/9-flex-tdTomato (Boston Children’s Hospital Vector Core, titer 1.5-2.5×10^13^ genome copies/ml) was inserted into the vagal complex. Virus solution was expelled using a Nanoject II injector (Drummond). Animals recovered from surgery were returned to housing. Mice were sacrificed 3 weeks later for the VG, the lung and the brainstem harvest.

#### Tissue collection and immunofluorescence staining

Mice were euthanized by CO_2_ inhalation. VG were dissected and harvested first. Then mice were then transcardinally perfused with PBS followed by perfusion fixation with ice-cold 4% paraformaldehyde (PFA). The lungs were then inflated with 4% PFA at 35cm H_2_O airway pressure and harvested. Lastly, the brainstems were harvested. All the tissues were fixed overnight at 4°C in 4% PFA.

After fixing, tissues were washed three times in PBS. Tissues were then cleared by incubation in CUBIC R1 solution on a rotating shaker (50rpm) at 4°C for 2-3 weeks until visually cleared.

For sectional analysis of fluorescent signals in the medulla, fixed brainstems were washed in PBS to remove residual PFA and then cryo-embedded in OCT compound following standard procedures. Blocks were sectioned at 40um in rostral to caudal sequence and screened for fluorescence using a Leica SP8 confocal microscope.

For immunostaining, cryo-embedded lung blocks were sectioned at 99um per slice and processed for immunostaining following a standard protocol. Primary antibodies used include mouse anti-alpha Smooth Muscle Actin (Sigma), Syrian hamster anti-PDPN (DSHB), rabbit anti-Von Willebrand Factor (Sigma), rabbit anti-Von Willebrand Factor (Sigma), rabbit anti-LYVE-1 (Abcam), rabbit anti-TAGLN (SM22, Abcam), rat anti-AGER (RAGE, R&D), rabbit-anti pro-SPC (Millipore) and rabbit anti-CGRP (Sigma). Secondary antibodies used include goat anti-rabbit FITC, goat anti-Syrian hamster FITC, goat anti-rat FITC, goat anti-rabbit Cy5 (all from Jackson ImmunoResearch Labs). Slides were mounted using Vectashield (Vector Labs) and imaged using a Leica SP8 or Nikon A1 confocal microscope.

#### Light sheet fluorescence microscopy

CUBIC buffers were prepared according to Susaki *et al*^34^. After sufficient tissue clearing in R1 buffer, tissues were embedded in 2% low melting agarose and then incubated in R2 solution at room temperature overnight prior to imaging. Cleared samples were imaged using a Zeiss Z.1 light sheet fluorescence microscope (LSFM). VG samples were imaged using a 5x objective (LSFM 5x NA 0.1) and a 1.45 5X Clarity chamber. Lung and brainstem samples were imaged using a 2.5x objective (LSFM 2.5x NA 0.1) and a 1.45 2.5X Clarity specific chamber (Translucence Biosystems).

#### Vagal RNAscope *in situ* hybridization

To prepare sections for RNAscope, fresh VG were frozen in OCT compound. Thin sections (12um) were prepared using a cryostat, collected on Superfrost Plus slides, and left to dry in the cryostat for at least 30min before staining. All staining procedures were performed using RNAscope Fluorescent Multiplex Kit (Advanced Cell Diagnostics, 320850) following the manufacturer’s instruction. Briefly, sections were post-fixed in 4% PFA for 1hr at room temperature (RT), washed in PBS, dehydrated in a series of ethanol washes, and then dried. A hydrophobic barrier was drawn around the section with an ImmEdge pen (Vector Lab, H-4000). The sections were treated with Protease IV in a HybEZ Humidity Control Tray for 30min at room temperature, incubated with target probes in a HybEZ Oven for 2hrs at 40°C, and then treated with Hybridize Amp. Slides were mounted using fluoromount-G with DAPI (Southern Biotech).

## Acknowledgments

We would like to thank members of the Sun lab, Drs. Axel Nimmerjahn, Seung Han (Salk), Drs. Binhai Zheng, Takaki Komiyama and Byungkook Lim (UCSD), Dr. Fan Wang (Duke), and Dr. Thomas Taylor-Clark (University of Southern Florida) for inputs and discussions. We thank Dr. Darrell Kotton and Dr. Laertes Ikonomou for sharing the *Nkx2-1^GFP^* mice, Jennifer Santini and Marcella Erb at the UCSD Light Microscopy Core and Eric Griffis at the UCSD Nikon Imaging Center for assistance with imaging and analysis. This work was supported by NIH SPARC OT2OD023857 (to X.S) and AHA 19POST34450103 (to Y.S.) and NHLBI F32 HL151168 (to J.B.).

## Author contributions

The authors confirm contribution to the study as follows: study conception and design: Y.S., J.B., X.S.; data collection: Y.S., J.B.; analysis and interpretation of results: Y.S., J.B., A.J., J.X., J.V.; draft manuscript preparation: Y.S., J.B., X.S. All authors reviewed the results and approved the final version of the manuscript.

