## Supplemental Files for "Mapping the Central and Peripheral Projections of Lung Innervating Sensory Neurons"

Su et al.
Supplementary Fig.1

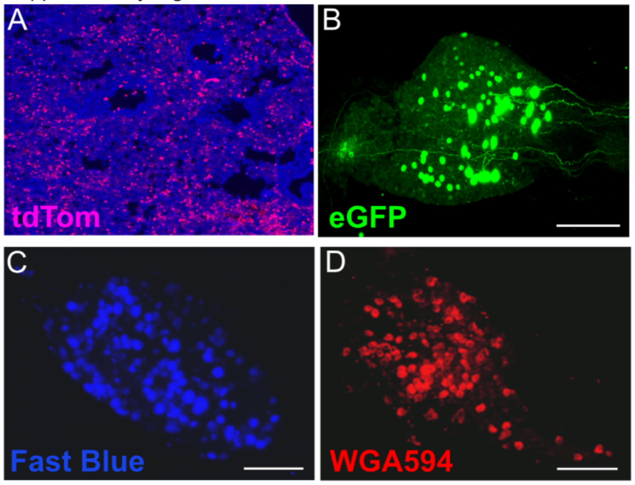


**Supplementary Fig.1. Retrograde tracing of lung innervating afferents led to labeling of cell bodies in the vagal ganglia.**
**(A)** Representative section of *Rosa-lxl-tdTomato* (Ai14) lung following intratracheal instillation of rAAV2-retro-cre, showing widespread infection. **(B-D)** Vagal ganglia from retrograde labeling using rAAV2-retro-eGFP (eGFP, B), Fast Blue (C), and wheat germ agglutinin (WGA594, D) instilled in wild-type lung. All scale bars 200um.

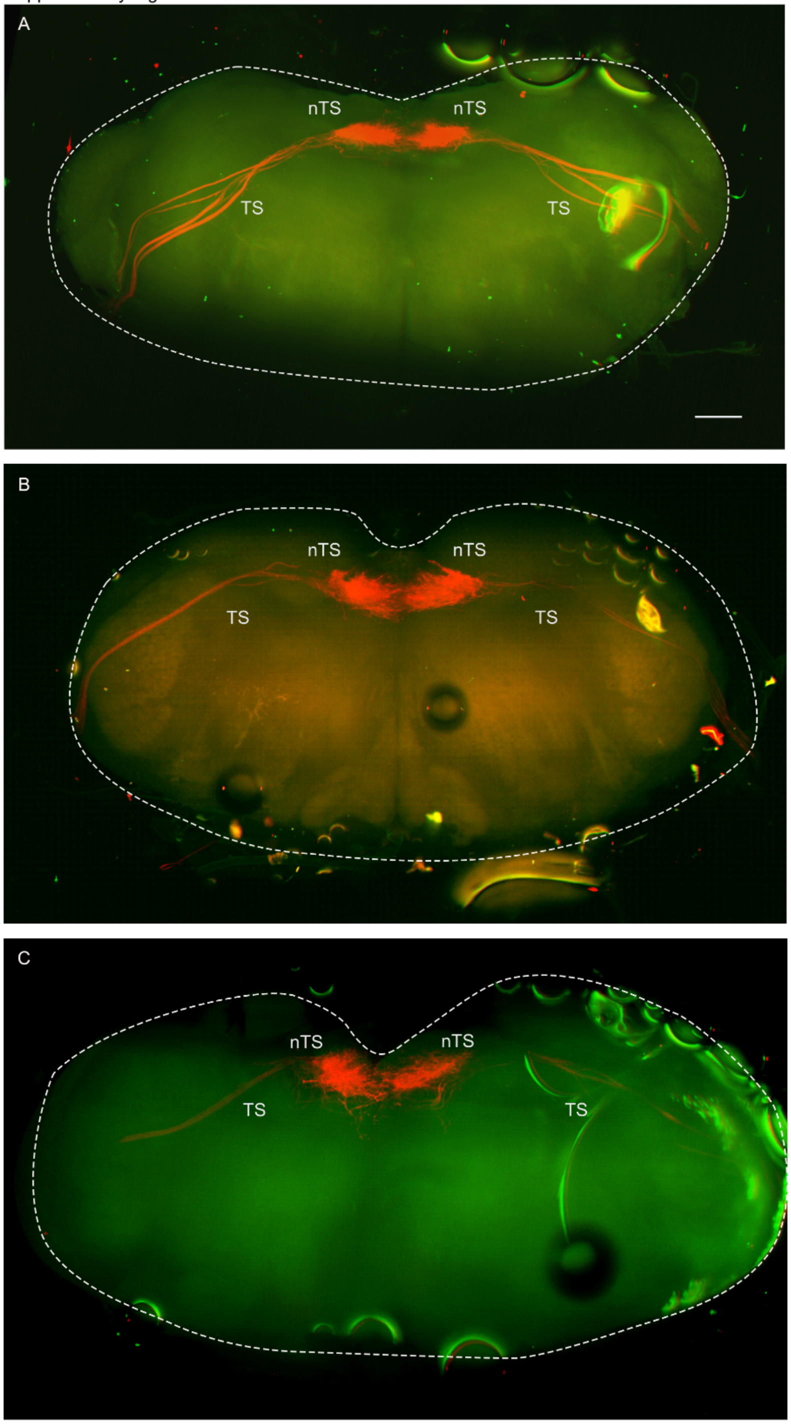


**Supplementary Fig.2. Retrograde tracing of lung innervating afferents densely arborized in the nTS bilaterally.**

**(A-C)** CUBIC cleared whole brainstem showing similar nerve terminal signals in bilateral nTS from another three of *Ai14* mice with intratracheal instillation of rAAV2-retro-cre. Lung-innervating neurons centrally project to the nTS via the TS. The intensity of TS signals varied based on exposure time set primarily based on the intensity of nTS signals. Dashed circles outline the brainstem. nTS, nucleus of the solitary tract; TS, solitary tract. Scale bar 500um.

Su et al.
Supplementary Fig.3

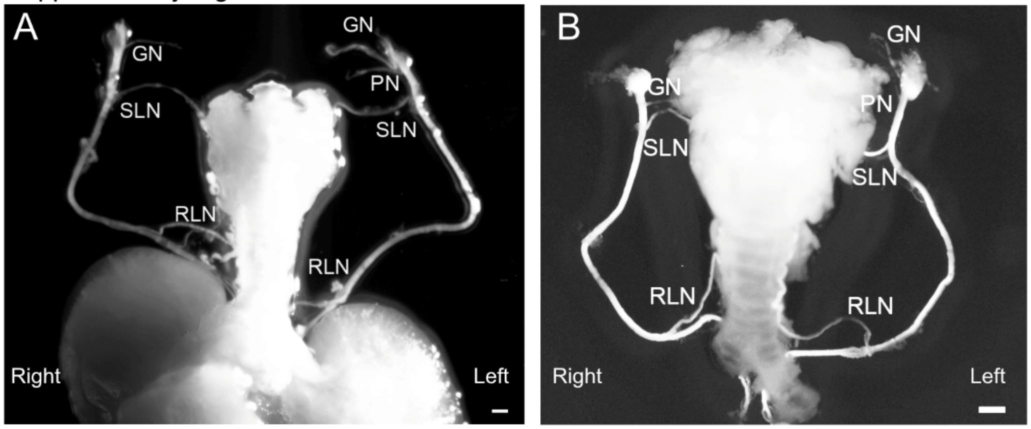


**Supplementary Fig.3. Sensory nerve paths that innervate the respiratory tract.**
**(A)** Bright field of blunt dissection from the *Vglut2-Cre; Ai14* mouse in Fig2.C-G, showing vagal nerve patterns. **(B)** Similar innervation patterns from another *Vglut2-Cre; Ai14* mouse in bright field. GN, glossopharyngeal nerve; PN, pharyngeal nerve; SLN, the superior laryngeal nerve; RLN, the recurrent laryngeal nerve. All scale bars 1mm.

Su et al.
Supplementary Fig.4

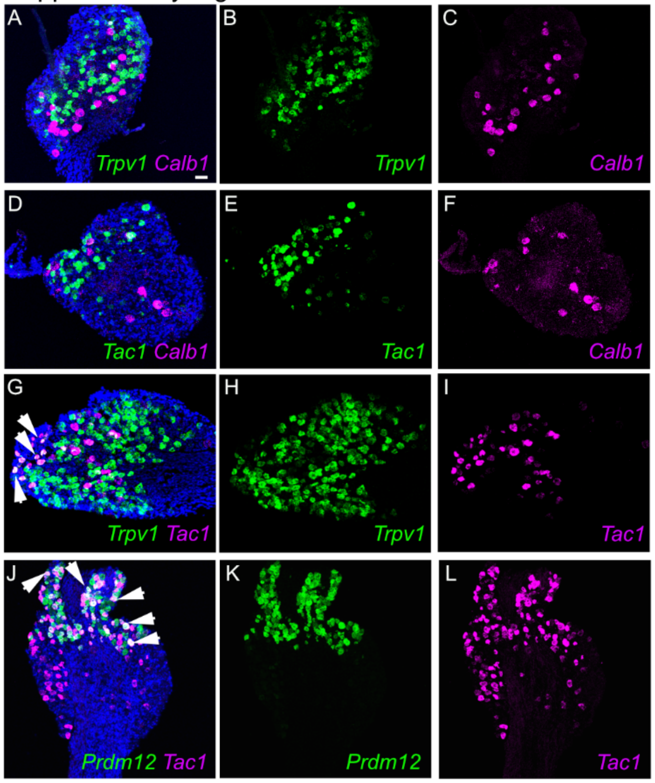


**Supplementary Fig.4. Vagal ganglia RNAscope showing gene expression overlap among sensory neuron markers.**
**(A-I)** Two color RNAscope reveals little to no overlap between *Calb1* with either *Trpv1* or *Tac1* vagal neurons, and some overlap between *Trpv1* and *Tac1*. **(J-L)** Two color RNAscope reveals *Tac1* expression in both jugular (*Prdm12*+) and nodose (*Prdm12*-) ganglia. Arrowheads point to neurons with double staining. Scale bar 50um.

Su et al.
Supplementary Fig.5

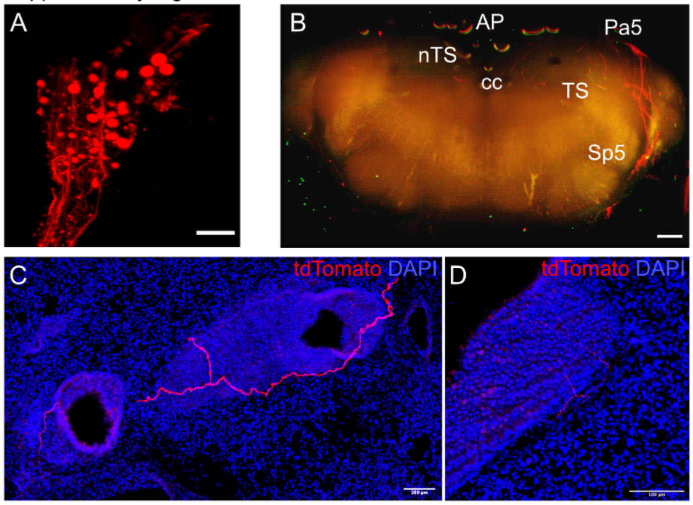


**Supplemental Fig.5. Brainstem and lung projection of *Piezo2-cre* mouse after unilateral vagal ganglia injection of AAV-flex-tdTomato.**
**(A, B)** Representative CUBIC cleared vagal ganglia (A) and whole brainstem (B) after AAV-flex-tdTomato vagal microinjection to the *Piezo2-cre* mouse. Autofluorescence green channel as background to outline the brainstem (B). nTS, the nucleus of solitary tract; TS, the solitary tract; AP, the area postrema; Pa5, the paratrigeminal nucleus; Sp5, the spinal trigeminal nucleus, cc, central canal. **(C, D)** Sections (99um) of the lung showing few fibers near airways. Scale bars 200um in (A), 500um in (B), 100um in (C, D).

Su et al.
Supplementary Fig.6

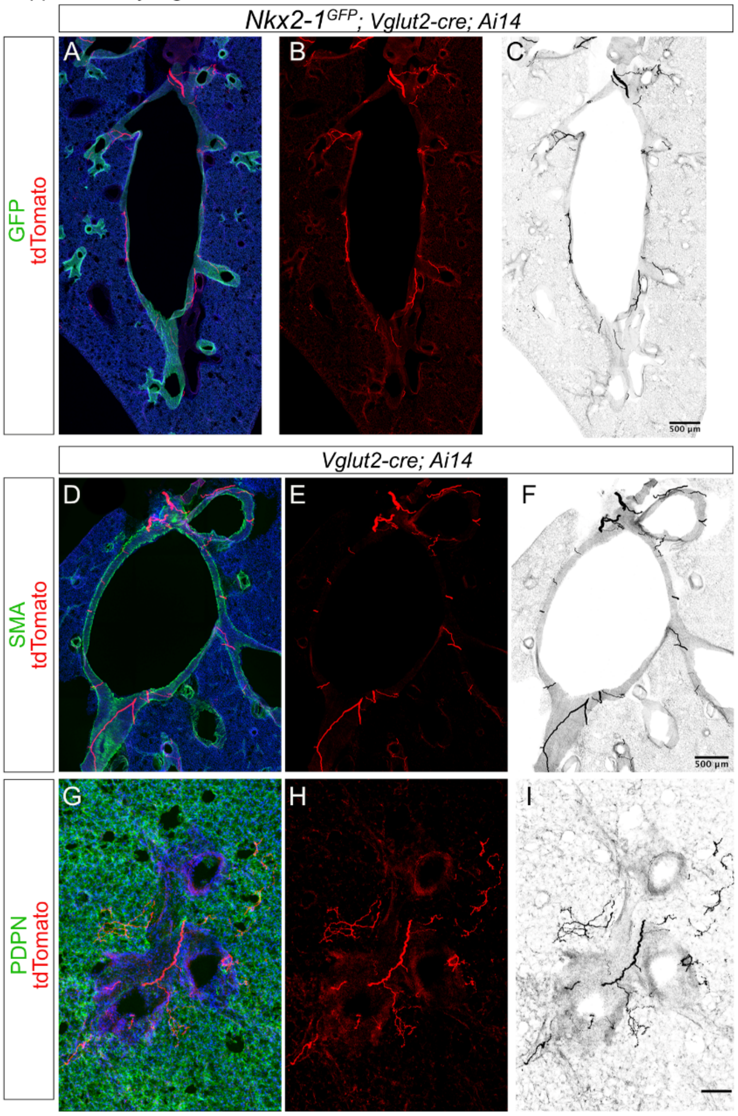


**Supplemental Fig.6. Innervation of airways by *Vglut2-cre; Ai14* nerve fibers.**
**(A-C)** Sections (99um) of the lung showing thick nerve bundles expressing tdTomato driven by *Vglut2-cre* project along large airways in which epithelial cells express GFP driven by *Nkx2-1* transcriptional control. **(D-F)** Thick nerve bundles expressing tdTomato driven by *Vglut2-cre* project along large airways and are localized near airway smooth muscle (SMA). **(G-I)** Nerve fibers expressing tdTomato (H-I) driven by *Vglut2-cre* project to smaller, distal airways (bronchioles) and the alveolar region (PDPN). Scale bars 500um in (C, F), 100um in (I).

Su et al.
Supplementary Fig.7

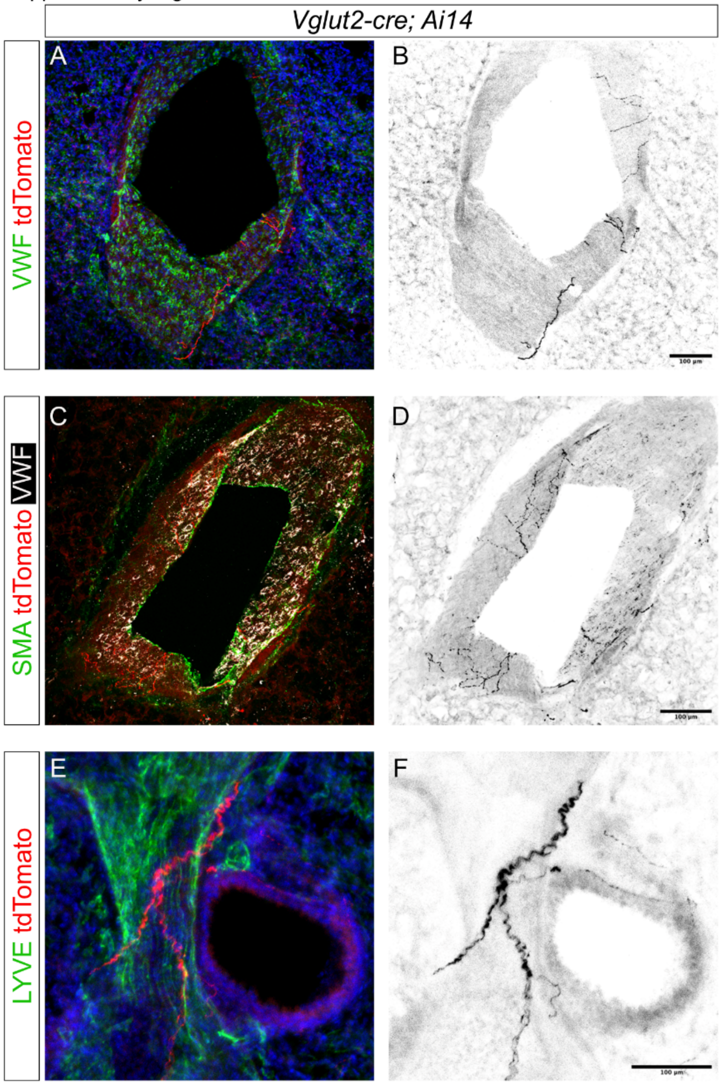


**Supplemental Fig.7. Innervation of vasculature by *Vglut2-cre; Ai14* nerve fibers.**
**(A, B)** Sections (99um) of the lung showing thin nerve fibers expressing tdTomato (red A; black B) driven by *Vglu2-cre* project to VWF expressing vasculature. **(C, D)** Thin nerve fibers expressing tdTomato driven by *Vglut2-cre* project to VWF expressing vasculature associated with vascular smooth muscle (SMA). **(E, F)** Nerve fibers expressing tdTomato driven by *Vglu2-cre* project to lymphatics expressing LYVE, near an airway. All scale bars 100um.

Su et al.
Supplementary Fig.8

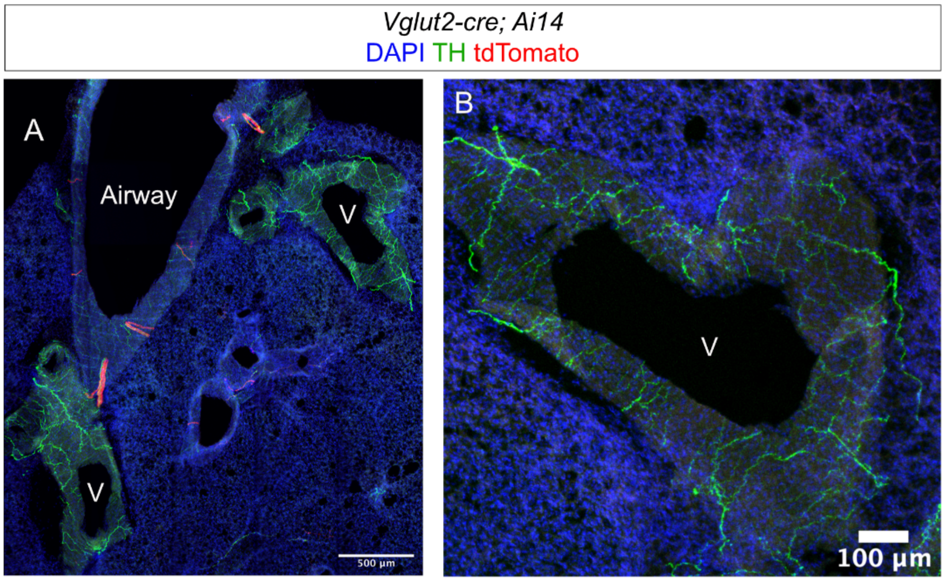


**Supplemental Fig.8. Dense innervation of vasculature by sympathetic nerve fibers expressing tyrosine hydroxylase.**
**(A, B)** Sections (99um) of the lung showing thick bundled nerve fibers expressing tdTomato driven by *Vglu2-cre* project to proximal airways. A few tyrosine hydroxylase (TH) expressing nerve fibers were found near proximal airways (A) but more were found to densely innervate vasculature (V in A, B). Scale bars 500um in (A) and 100um in (B).

Su et al.
Supplementary Fig.9

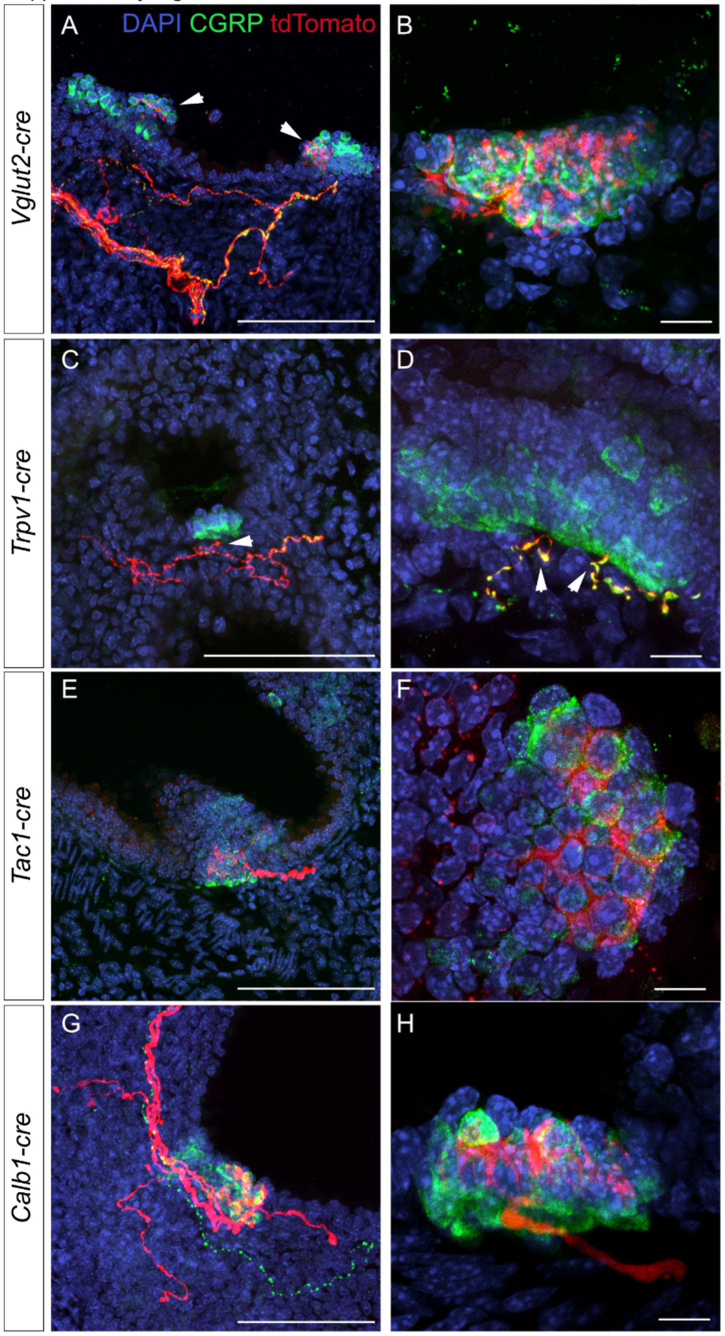


**Supplemental Fig.9. Vagal sensory neuron innervation of PNECs.**
**(A-H**) Sections (99um) of the lung showing vagal nerve projections innervating PNECs following vagal injection of *AAV-flex-tdTomato* into respective lines as labeled. Arrowheads in A point to innervated clusters. *Trpv1-cre* nerve fibers project to the basal side of PNECs (C, D, arrowheads), and these nerve fibers can be CGRP+ (D). *Tac1-cre* (E, F) and *Calb1-cre* (G, H) nerve fibers intercalate within the PNEC cluster. Scale bars 100um in (A, C, E, G), 10um in (B, D, F, H).

**Supplemental movie 1. Light sheet movie of *Rosa-lxl-tdTom (Ai14)* vagal ganglia and nerve tracts following intratracheal instillation of rAAV-retro-cre and CUBIC clearing (Fig.1C).**

**Supplemental movie 2. Light sheet movie of *Vglut2-cre; Ai14* CUBIC cleared right cranial lung lobe (Fig.2I).**

**Supplemental move 3. Light sheet movie of *Nkx2-1*$^{GFP}$*; Vglut2-cre; Ai14* CUBIC cleared right cranial lung lobe (Fig.2K).**

**Supplemental movie 4. Innervation of PNECs by vagal injection of AAV-flex-tdTomato into *Vglut2-cre* line (Fig.9B).**

**Supplemental movie 5. Innervation of PNECs by vagal injection of AAV-flex-tdTomato into *Trpv1-cre* line (Fig.9E).**

**Supplemental movie 6. Innervation of PNECs by vagal injection of AAV-flex-tdTomato into *Tac1-cre* line (Fig.9H).**

**Supplemental movie 7. Innervation of PNECs by vagal injection of AAV-flex-tdTomato into *Calb1-cre* line (Fig.9K).**
